# FloREN: Decoding Immune Regulatory Networks through Interpretable Graph Transformer Patient Representations

**DOI:** 10.64898/2026.07.12.738088

**Authors:** Iñigo Clemente-Larramendi, Sophie Hillion, Divi Cornec, Christophe Jamin, Nathan Foulquier

## Abstract

Single-cell RNA sequencing (scRNA-seq) enables detailed characterization of cellular heterogeneity, yet understanding the full cellular and regulatory environment of complex tissues remains challenging. In the era of large single-cell atlases, this technology has become increasingly accessible, and datasets have grown in scale and statistical power. As a result, sample representation methods have emerged as a promising strategy to summarize patient-level biological variation. However, most existing approaches rely on unsupervised learning frameworks with ambiguous biological interpretability.

Here we present a Framework for Learning Over REgulatory-Embedding Networks (FloREN), a supervised and interpretable sample representation method. FloREN models single-cell data as a heterogeneous network integrating cells and genes together with gene regulatory and cell-cell communication relationships. Through condition-aware embeddings and interpretable attention networks, FloREN enables improved sample stratification and biomarker discovery. In addition, the framework supports downstream analyses that found specific immune network mechanisms in immune-mediated inflammatory diseases (IMIDs).

## Introduction

Advances in high-throughput technologies over the past decade have enabled the generation of increasingly large and complex biological datasets, transforming our ability to study human disease at unprecedented resolution^1–4^. Among these technologies, single-cell RNA sequencing (scRNA-seq) has emerged as a cornerstone for dissecting cellular heterogeneity. Profiling hundreds of millions of cells across diverse tissues and disease contexts has revealed novel cell types^5^, context-specific marker genes^6,7^, gene programs linked to therapeutic response, and clinically relevant patient subpopulations^8,9^. As scRNA-seq studies expand toward cohort-scale analyses, methods that summarize each sample as a single vector, commonly termed sample representation methods, have become increasingly important. These approaches capture coordinated changes across tissue ecosystems and enable direct linkage between sample-level phenotypes and their underlying cellular and molecular composition^10,11^.

Existing methods can be broadly divided into unsupervised and supervised approaches. Unsupervised methods aim to capture global similarities between samples without relying on labeled outcomes, and are effective for sample stratification, batch correction, and exploratory analysis. Supervised methods instead learn embeddings optimized for downstream tasks such as disease classification or outcome prediction, where interpretability becomes essential for translating learned representations into candidate biomarkers or drug targets.

The simplest strategy is pseudobulk aggregation, which averages gene expression across cells within each sample. Although pseudobulk profiles retain substantial biological information and support accurate patient classification^12^, they are difficult to relate back to cell-type-specific or single-cell-resolved patterns. Compositional approaches instead represent samples through the distribution of cell states: GloScope^13^ models each sample as a distribution over cell annotations using Gaussian mixture models (GMMs) on PCA embeddings in an unsupervised setting, while CloudPred^14^ trains a supervised classifier on similar GMM-based representations; both, however, remain difficult to interpret at the level of individual genes or pathways. An alternative strategy represents each sample using a broad panel of precomputed single-cell statistics (differential abundance (DA), differential gene expression (DGE), gene set enrichment analysis (GSEA), and cell-cell communication scores) as implemented in scFeatures^15^. More recently, the machine learning community has developed foundation models for single-cell cohorts, including PaSCient^11^ and PULSAR^17^, which aim to disentangle biological from technical variation in an unsupervised setting, and mcBERT^16^, which leverages large-scale pretraining to improve supervised predictive performance; several of these incorporate learned attention weights in an effort to improve interpretability of cell- and gene-level contributions. SampleCLR^18^ combines contrastive self-supervised and supervised objectives over PCA embeddings and outperforms earlier foundation models showing that the potential of this techniques, though its interpretability remains limited to cell-level weights. MrVI^10^, which relies on an unsupervised variational inference framework, extends interpretability further by linking sample representations to DGE and DA analyses. Collectively, these approaches reveal a persistent trade-off: no existing method combines a clearly supervised objective with interpretability across multiple biological levels: cells, genes, and their interactions.

To address this gap, we introduce FloREN (Framework for Learning Over REgulatory-Embedding Networks), a graph neural network (GNN)-based method that models sample-level single-cell data as biological networks and leverages attention-based interpretability to advance the state of the art in sample representation learning.

Biological systems are inherently networked, with genes, proteins, and cells interacting in coordinated ways; these relationships can be naturally represented as graphs, in which nodes denote biological entities and edges encode their interactions^19,20^. As datasets grow in scale and complexity, graph-based modeling has emerged as a powerful framework for integrating and analyzing high-dimensional biological data^20^. Accordingly, GNNs have been widely adopted in single-cell analysis, enabling applications such as data imputation^21–25^, clustering^26–32^, and cell-type annotation^33–38^. Despite this progress, most GNN-based approaches rely on homogeneous graphs that typically represent only cells and their transcriptomic similarity^39^, overlooking the inherent heterogeneity of biological systems, a limitation that is especially consequential for sample-level inference. Only a handful of methods move beyond this paradigm: DeepMAPS^40^ and Segger^41^ incorporate heterogeneous graphs but are designed for clustering and segmentation tasks, respectively, while Cellograph^42^ and OmicsGAT^43^ use more limited graph structures for differential expression or outcome prediction. There thus remains a clear need for models that fully exploit biological heterogeneity to generate interpretable, patient-level representations.

This challenge is especially relevant to immune-mediated inflammatory diseases (IMIDs), where pathology arises from system-wide dysregulation of the immune response^44^. These diseases, including rheumatoid arthritis (RA), systemic lupus erythematosus (SLE), and multiple sclerosis (MS), involve complex interactions among diverse immune cell populations, and their pathology reflects coordinated network-level alterations rather than isolated cellular changes, making IMIDs a stringent benchmark for evaluating sample representation methods^45^.

That’s why we propose FloREN, an interpretable sample representation framework that models single-cell data as heterogeneous biological networks. From SampleCLR contrastive self-supervised and supervised promising combination, FloREN combines heterogeneous graph transformers, graph self-supervised learning, and supervised optimization to generate biologically informed patient embeddings while preserving interpretability through attention-based prioritization of cells, genes, and regulatory interactions. By jointly integrating cellular composition, gene regulation, and intercellular communication within a unified graph representation, FloREN enables both accurate patient stratification and mechanistic discovery from single-cell datasets.

## Results

### The architecture of FloREN

FloREN is a graph neural network (GNN) framework designed to extract biologically meaningful patterns from scRNA-seq data by jointly modeling multi-layered relationships within heterogeneous graphs, gene expression, regulatory interactions, and intercellular communication (**Fig. 1a**). FloREN takes as input a single-cell expression object with log-normalized counts, restricted to a defined gene set for the analysis (typically highly variable genes, HVGs). Beyond gene expression, FloREN can also operate directly on transcription factor (TF) activity scores derived from tools such as DecoupleR^63^, allowing the framework to model regulatory state rather than expression alone when desired. FloREN first projects cells and genes into a shared latent space using paired autoencoders. This denoising step generates comparable low-dimensional representations that enable robust inference of heterogeneous biological interactions while reducing the impact of sparse single-cell measurements.

**Fig. 1.**
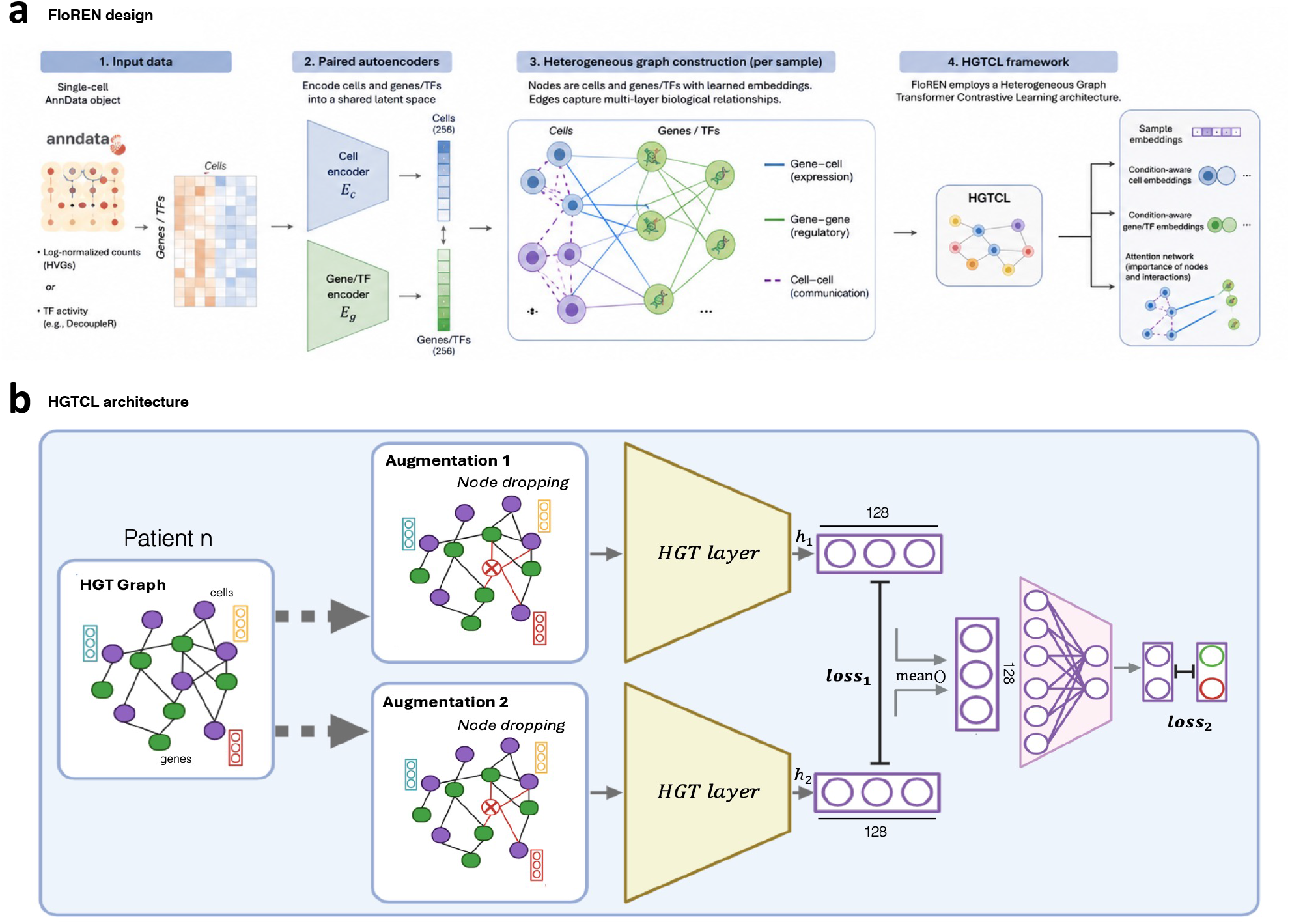
Overview of the FloREN architecture. **a**, The overall FloREN pipeline. Starting from scRNA-seq normalized expression matrices, the workflow includes six major steps and two learning layers: an Autoencoder (AE) for dimensionality reduction and the novel Heterogenous Graph Transformer Contrastive Learning (HGTCL) module for sample representation learning. Graph construction incorporates biological priors such as gene expression, transcription factor activity, cell-cell communication (via *CellChat*), and gene-gene correlations (via a Bayesian scoring scheme combining correlations and prior knowledge). **b**, Illustration of the HGTCL classifier process for extracting sample-level embeddings. From each sample graph, two augmentations are generated by node dropping. Both pass through a HGT encoder, and their mean-pooled node embeddings are used as graph-level representations. These embeddings are trained to both (1) maximize agreement between augmentations (contrastive loss), and (2) classify sample labels (supervised loss), enabling robust and interpretable feature extraction.

The core of FloREN is to use this same dimensionality space to construct a heterogeneous graph for each sample, in which nodes correspond to cells and genes (or TFs), and node features correspond to their learned embeddings. Edges are defined across three biologically motivated interaction layers. First, gene-cell edges link each gene to the cells expressing it. Second, gene-gene edges capture regulatory relationships inferred through a Bayesian scoring strategy (see **Methods**) that integrates curated immune interaction databases with correlation estimates, enabling detection of both known and novel regulatory associations. Third, optional cell-cell communication edges can be incorporated using curated ligand-receptor interaction networks, for example via CellChat^66^. This modular design gives FloREN substantial flexibility in graph construction, allowing users to include or omit interaction layers depending on data availability and biological question.

These heterogeneous graphs are processed using our Heterogeneous Graph Transformer Contrastive Learning (HGTCL) architecture (**Fig. 1b**) which combines the Heterogeneous Graph Transformer (HGT)^71^ with graph Contrastive Learning (graphCL)^67^. Because biological graphs derived from single-cell data are inherently noisy and sensitive to graph construction, HGTCL first learns robust sample representations in a self-supervised manner by maximizing agreement between two stochastic augmentations of the same graph generated through node dropping. Within each HGT layer, relation-specific attention propagates information across cells, genes, and biological interactions, enabling context-aware integration of heterogeneous signals while remaining invariant to minor perturbations of the graph. Following the HGT layers, node embeddings are aggregated by mean pooling into a single fixed-dimensional sample representation. Although contrastive learning produces robust graph representations, it does not ensure that they capture disease-relevant variation. FloREN therefore introduces a supervised multilayer perceptron (MLP) operating on the learned sample embeddings. This objective steer representation learning toward phenotype-associated biological differences while preserving the structural information captured during self-supervised training, enabling both multitask and multiclass prediction (see **Methods**)^46,47^.

Together, these components allow FloREN to produce, from a single training pass, three complementary outputs: supervised sample-level embeddings from the MLP classifier, condition-aware cell, and gene embeddings from the HGT encoder, and an interpretable attention network describing cell-, gene-, and interaction-level contributions to each sample’s representation, enabling a range of downstream analyses discussed below.

### FloREN demonstrates superior performance across diverse single-cell studies

To evaluate FloREN’s generalizability, we applied it to five independent IMIDs scRNA-seq datasets spanning diverse immunological contexts, experimental designs, and patient cohorts (**Fig. 2a**; see **Methods**)^48–52^. For each dataset, we constructed patient-level heterogeneous graphs reflecting the specific structure of that study; together, these five scenarios also serve as an ablation test of FloREN’s performance across varying graph compositions and data regimes.

**Fig. 2.**
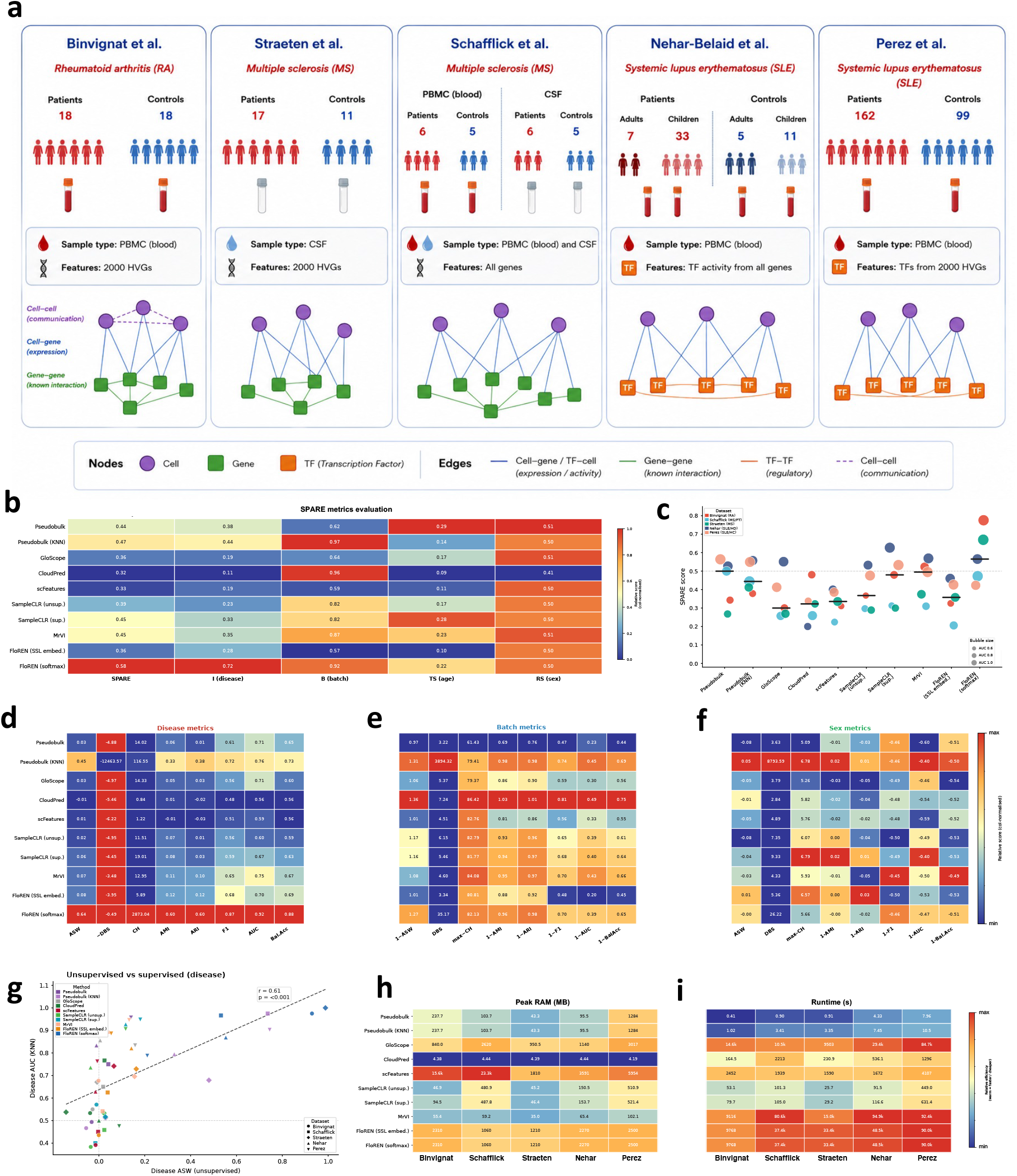
FloREN improves predictive performance while comparable computational effort. **a**, Overview of the five independent datasets used to evaluate FloREN, covering diverse sample sources (blood, CSF, or both), disease states (rheumatoid arthritis RA, multiple sclerosis MS, systemic lupus erythematosus SLE, control CNT), and graph construction approaches **b**, Comparison of SPARE metrics between sample pseudobulk (“Pseudobulk”), kNN classifier on the pseudobulk embeddings (“Pseudobulk (KNN)”), GloScope, Cloudpred, scFeatures, unsupervised (“SampleCLR (unsup.)”) and supervised (“SampleCLR (sup.)”) SampleCLR, MrVI, FloREN’s self-supervised (“FloREN (SSL embed.)”) and final classfier (“FloREN (softmax)”) embeddings across aggregated score (“SPARE”), disease information retention (“I (disease)”), batch information removal (“B (batch)”), age trajectory conservation (“TS (age)”) and sex robustness (“RS (sex)”). **c**, Aggregated SPARE score across datasets for each method. **d**, Extended disease information retention for each method based in unsupervised clustering metrics (Average Silhouette Width (ASW), inverse Davies-Bouldin Score (−DBS), and Calinski-Harabasz Index (CH)), supervised clustering metrics (Adjusted Mutual Information (AMI) and Adjusted Rand Index (ARI)), and predictive performance metrics (5-fold cross-validation F1 score (F1), Area Under the Receiver Operating Characteristic Curve (AUC), and Balanced Accuracy (BalAcc)). Higher the better. **e**, Same extended analysis for each method but on batch information removal (inverse of the original scores). Higher the better. **f**, Same extended analysis for each method but on sex information removal (inverse of the original scores). Higher the better. **g**, Correlation between the disease AUC score and the disease ASW for every method and every dataset. **h**, Peak RAM used in megabytes (MB) per method per dataset. **i**, Time used in second (s) per method per dataset.

We benchmarked FloREN against a panel of leading, widely used methods spanning the main strategies described above. As baselines for gene-expression information retention, we used sample-level pseudobulk profiles and a k-nearest-neighbors (kNN) classifier applied to pseudobulk embeddings. As baselines for cell-composition information retention, we used GloScope, which models a Gaussian mixture model (GMM) over cells’ PCA embeddings in an unsupervised setting, and its supervised counterpart CloudPred, which fits a patient classifier using the same GMM-based representation. We additionally included scFeatures, an algorithmic approach that computes a comprehensive panel of single-cell summary statistics per patient, to assess whether learned representations capture information beyond what these hand-crafted statistics already provide. We further benchmarked against both unsupervised and supervised variants of SampleCLR; making us exclude PULSAR, PaSCient, or mcBERT, as these were benchmarked and proved substantially more difficult to adapt to our datasets. Finally, we included MrVI as a representative of variational-inference-based single-cell (scVI-family) approaches, an increasingly prominent strategy in this field. All baselines were compared against two FloREN outputs: the self-supervised contrastive embeddings, and the final layer of the supervised FloREN classifier.

Across all five datasets, FloREN embeddings outperformed alternative sample-level representation methods according to the Sample Preservation and Representation Evaluation (SPARE) metrics^53^(see **Methods**). FloREN achieved the highest overall SPARE score (**Fig. 2b**), driven primarily by superior disease-information retention, while remaining competitive in batch-effect removal, age-trajectory conservation, and robustness to sex. Self-supervised FloREN embeddings performed comparably to other unsupervised methods, including unsupervised SampleCLR, scFeatures, and GloScope, but fell behind MrVI in this setting (**Fig. 2c**). In contrast, the supervised FloREN classifier’s final-layer embeddings consistently outperformed all supervised and unsupervised baselines across SPARE metrics. The FloREN classifier’s advantage was most pronounced for disease-information retention, where it obtained the highest scores across unsupervised clustering metrics (Average Silhouette Width [ASW], inverse Davies-Bouldin score [−DBS], and Calinski-Harabasz index [CH]), supervised clustering metrics (Adjusted Mutual Information [AMI] and Adjusted Rand Index [ARI]), and predictive-performance metrics (5-fold cross-validated F1 score, area under the receiver operating characteristic curve [AUC], and balanced accuracy) (**Fig. 2d**). For batch- and sex-information removal, FloREN’s performance was comparable to the top baselines, with the kNN classifier on pseudobulk embeddings and CloudPred achieving the best scores on these specific tasks (**Fig. 2e,f**). The combined evaluation of supervised and unsupervised disease metrics, together with the agreement between SPARE and AUC scores, confirms the robustness of these results (**Fig. 2g**).

Finally, in terms of computational performance, FloREN required longer runtimes than PCA-based methods such as SampleCLR but was comparable to more computationally demanding approaches such as GloScope (GMM fitting) and MrVI (variational inference over the full count matrix) (**Fig. 2h**). FloREN’s memory usage was higher than that of SampleCLR or MrVI, but remained comparable to scFeatures, which computes thousands of statistics per patient (**Fig. 2i**).

### FloREN generates patient-level representations that reveal disease trajectories and intermediate SLE states

We used the Perez dataset^52^, originally designed to train supervised models distinguishing SLE patients from healthy donors, to evaluate whether supervised and unsupervised sample representation methods could additionally distinguish SLE patients experiencing disease flares (FLARE) from non-flaring SLE patients, a substantially higher disease-activity state.

We found this to be a considerably harder task: no clear separation between flare and non-flare status emerged in UMAP, t-SNE, PCA, or PHATE visualizations for any method, with the exception of GloScope, which we instead visualized using UMAP, MDS, Isomap, and spectral embedding, reflecting its distributional (GMM-based) representation. Despite the absence of clear visual separation, FloREN’s self-supervised embeddings ranked among the top-performing methods, alongside pseudobulk, the kNN classifier on pseudobulk embeddings, MrVI, and both unsupervised and supervised SampleCLR, based on 5-fold cross-validated F1, AUC, and balanced accuracy scores (**Fig. 3a**).

**Fig. 3.**
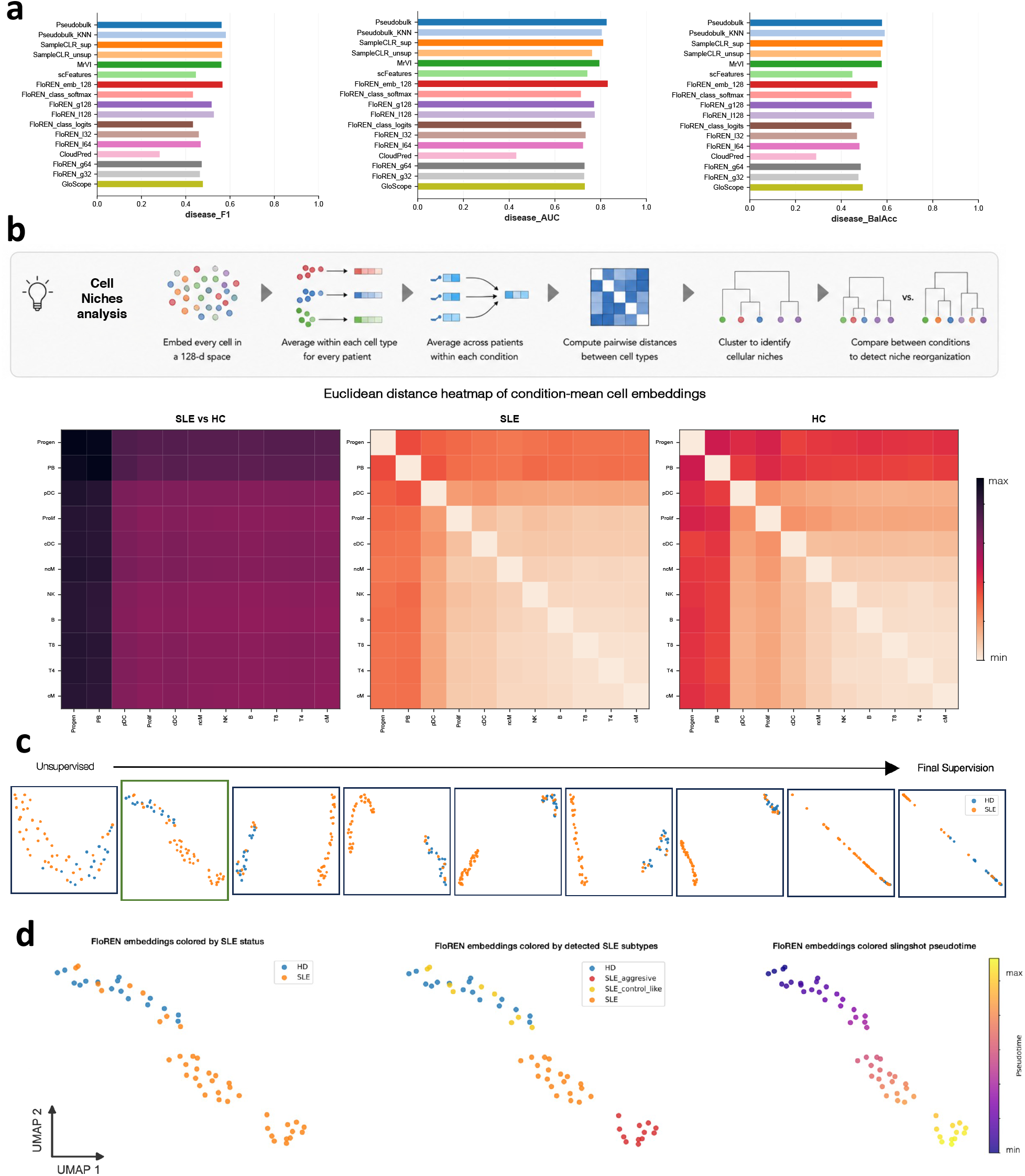
FloREN reveals disease trajectories and intermediate SLE states at the patient level. **a**, Predictive performance metrics (5-fold cross-validation F1 score (F1), Area Under the Receiver Operating Characteristic Curve (AUC), and Balanced Accuracy (BalAcc)) between methods towards HD, SLE and FLARE patients. Methods shown are sample pseudobulk (“Pseudobulk”), kNN classifier on the pseudobulk embeddings (“Pseudobulk (KNN)”), GloScope, Cloudpred, scFeatures, unsupervised (“SampleCLR (unsup.)”) and supervised (“SampleCLR (sup.)”) SampleCLR, MrVI and the different FloREN’s MLP layers: from FloREN’s self-supervised (“FloREN_embed_128)”), FloREN’s linear 128d layer (“FloREN_l128”), FloREN’s GELU activated 128d layer (“FloREN_g128”), FloREN’s linear 64d layer (“FloREN_l64”), FloREN’s GELU activated 64d layer (“FloREN_g64”), FloREN’s linear 32d layer (“FloREN_l32”), FloREN’s GELU activated 32d layer (“FloREN_g32”), FloREN’s linear 2d layer (“FloREN_class_logits”) and final classfier (“FloREN_class_softmax)”) **b**, Hierarchical clustering heatmaps based on Euclidean distances between condition-mean cell embeddings. **c**, UMAP visualization of sequential FloREN patient embeddings for the Nehar-Belaid dataset, showing the progression from unsupervised to fully supervised representation layers. **d**, UMAP visualization of the FloREN 128-dimensional patient embedding, colored by (1) original condition labels, (2) three SLE subgroups, an SLE-control-like cluster proximal to HDs, a canonical SLE cluster, and a distinct SLE-aggressive cluster, and (3) trajectory inference projected onto the embedding space using Slingshot.

Hierarchical clustering based on Euclidean distance between condition-mean cell embeddings showed that SLE and healthy donor (HD) samples were related but not fully aligned, with distinct cellular niches emerging within each condition (**Fig. 3b**). Plasmablasts and progenitor cells showed clear separation between SLE and HD samples, consistent with their high importance in the attention-based analysis. Within-condition clustering further identified a distinct activation substate specific to these two populations, alongside a broadly connected cluster comprising CD4+ and CD8+ T cells, classical and non-classical monocytes (cM/ncM), natural killer (NK) cells, and B cells. Proliferative cells, conventional dendritic cells (cDCs), and plasmacytoid dendritic cells (pDCs) formed a shared niche in HD samples, whereas pDCs separated from this niche in SLE patients, consistent with their established role as the primary source of type I interferon in autoimmune disease^54^.

To further assess whether FloREN embeddings capture continuous disease progression, we analyzed the Nehar-Belaid dataset^51^, in which patients are annotated with discrete SLE disease stages. FloREN distinguished SLE patients from healthy donors with high accuracy (F1 = 0.85, ROC-AUC = 0.80), and the position of intermediate-stage patients within the embedding space tracked with their annotated disease progression (**Fig. 3c**). A lower-dimensional projection of the FloREN embedding space further separated SLE patients from HDs (**Fig. 3d**) and revealed three distinct subgroups: SLE-control-like, canonical SLE, and SLE-aggressive. Trajectory inference using Slingshot^55^ recapitulated a continuous progression from HDs through these intermediate states to SLE-aggressive. This stratification was independent of batch, age, and sex effects.

### FloREN identifies disease-relevant cellular populations and gene signatures in multiple sclerosis

We next applied FloREN in a top-down analytical setting, using it as a first-line strategy for identifying key cellular and molecular drivers of disease directly from patient cohorts. We analyzed the Straeten dataset^49^, an integrated CSF and PBMC scRNA-seq compendium comprising 17 multiple sclerosis (MS) patients and 11 healthy donors (HD). The cell-type annotation performed by the original authors procedure^49,56^ yielded cell-type predictions at two levels of granularity: Level 1 (major cell types) and Level 2 (fine-grained subtypes).

Using the same dimensionality-reduction panel applied to the Perez dataset, we first confirmed that FloREN achieves the highest separation of MS patients from HDs among all benchmarked methods (**Fig. 4a**) and that this separation emerges progressively over the course of classifier training. We then computed the mean attention score for each cell type across patients to prioritize populations for downstream analysis. Monocytes emerged as the most influential cell type at Level 1 (**Fig. 4b**). At Level 2, CD14+ and CD16+ monocytes remained among the top-ranked populations, while AXL+SIGLEC6+ dendritic cells (ASDCs) showed the strongest attention signal. ASDCs represent a transitional population between plasmacytoid and conventional dendritic cells with strong T-cell-activating and migratory capacity^57,58^; their prominence in our analysis suggests a previously underappreciated role in shaping the neuroinflammatory environment in MS. FloREN also assigned high attention to several poorly defined clusters (“Un_assigned,” “other,” and “other_T”), suggesting that their original annotation may warrant reevaluation. Consistent with the attention-based prioritization, the relevance of key CD4+ subtypes was independently confirmed through UMAP visualization (**Fig. 4d**).

**Fig. 4.**
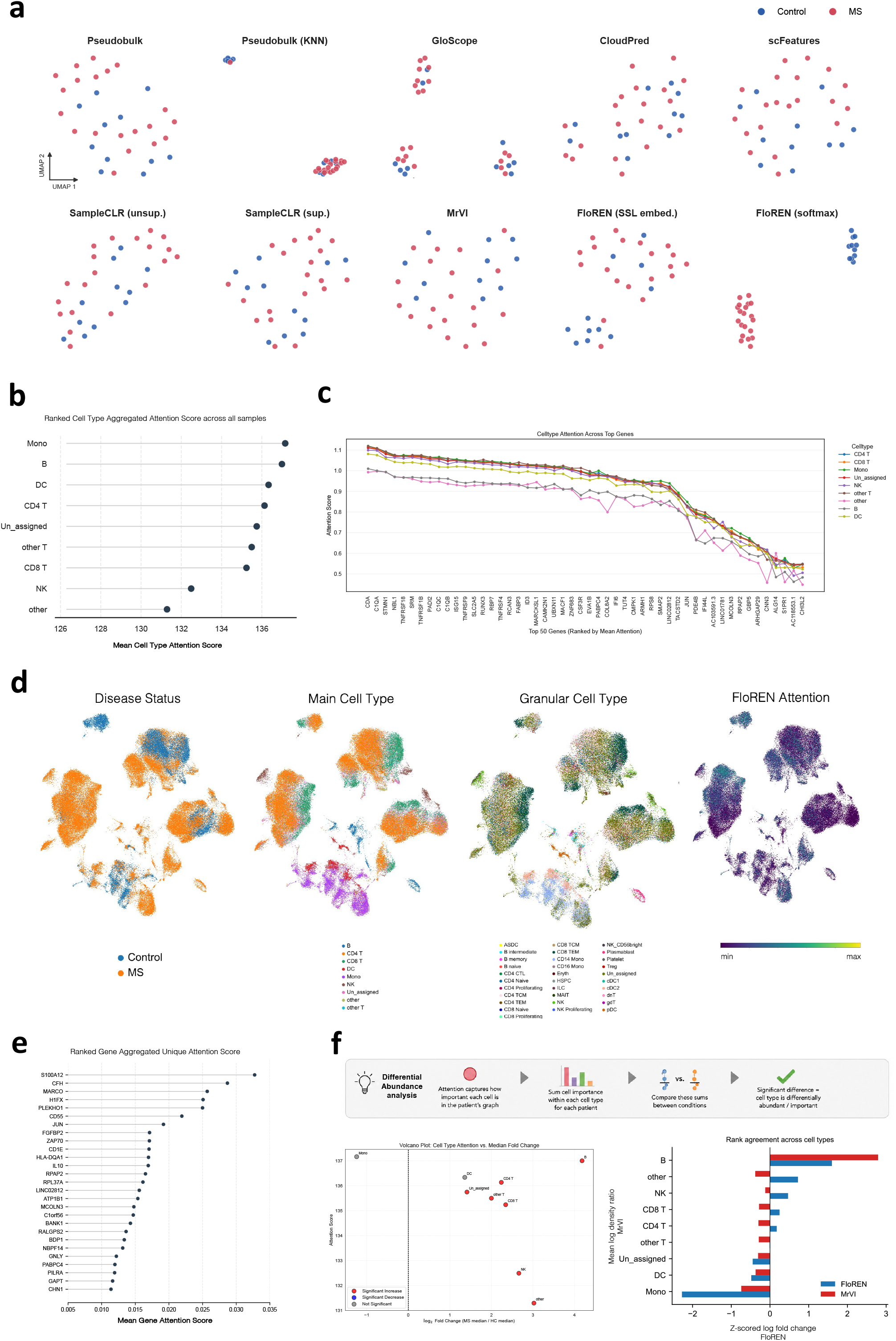
FloREN identifies disease-relevant cellular populations and gene signatures in multiple sclerosis. **a**, UMAP visualization of sample embeddings applied to Straeten dataset^45^ obtained from sample pseudobulk (“Pseudobulk”), kNN classifier on the pseudobulk embeddings (“Pseudobulk (KNN)”), GloScope, Cloudpred, scFeatures, unsupervised (“SampleCLR (unsup.)”) and supervised (“SampleCLR (sup.)”) SampleCLR, MrVI, FloREN’s self-supervised (“FloREN (SSL embed.)”) and final classfier (“FloREN (softmax)”). **b**, Bubble plot showing Level 1 cell types ranked by their mean attention score. **c**, Ranked genes according to their mean attention score across cell types. **d**, UMAP of cells colored by disease, Level 1 annotation, Level 2 annotation and FloREN attention. **e**, Bubble plot of genes ranked by their monocyte-enriched attention score. **f**, FloREN Differential Abundance (DA) analysis results volcano plot with log Fold Change and attention score. Agreement DA analysis between FloREN log Fold Change and MrVI Log Density Ratio.

We then examined gene-level attention within each cell type (**Fig. 4c**), which revealed a shared inflammatory signature spanning multiple immune populations: complement activation (C1QA/B/C), citrullination (PADI2), type I interferon signaling (ISG15, IFI6), costimulatory receptors (TNFRSF4, TNFRSF9, TNFRSF18), and tissue-residency factors (ZNF683, RUNX3). The prioritization of CHI3L2, a known MS biomarker^59^, further supports the biological relevance of FloREN’s attention-based rankings. To resolve cell-type-specific programs within monocytes, the most influential population identified above, we applied gene program decomposition (see **Methods**), uncovering a distinct monocyte-enriched program (**Fig. 4e**) centered on IL10 as a hub gene. Notably, this program also included markers typically associated with lymphoid lineages, such as ZAP70, FCRL5, and BANK1. We interpret this as more consistent with a hybrid activation state or strong intercellular communication signaling between myeloid and lymphoid compartments than with simple misannotation^49^, though we cannot fully exclude residual annotation noise. Additionally, the prominence of SATB1, A1CF, HLA-DRB1, and S100A12 suggests a broader molecular reprogramming of the myeloid lineage in MS.

Finally, we performed a differential abundance (DA) analysis using FloREN’s attention scores. Because FloREN assigns an attention score to each cell, cumulative attention per cell type can be computed for each patient; under the assumption of comparable per-cell attention contributions across conditions, a higher cumulative attention score for a given cell type reflects its increased relative representation in that condition. Using this approach, we observed increased attention-based representation of CD4^+^ T cells, CD8^+^ T cells, B cells, and NK cells in MS relative to HD (**Fig. 4f**). These findings agree with MrVI’s DA analysis for B cells, dendritic cells, and monocytes, but diverge for NK, CD4^+^, and CD8^+^ T cells (**Fig. 4f**).

### FloREN reveals regulatory gene programs and immune cell drivers of rheumatoid arthritis

We next applied FloREN as a bottom-up analytical framework, in contrast to the top-down approach used for MS, to the Binvignat dataset^48^, comprising PBMC samples from 18 rheumatoid arthritis (RA) patients and 18 matched healthy controls (HC). To capture intercellular interactions in this analysis, we incorporated CellChat-derived communication edges into the FloREN graph.

We confirmed that FloREN achieves the highest separation of RA patients from HCs among all benchmarked methods (**Fig. 5a**), with this separation emerging progressively over the course of classifier training. Attention-based ranking of cell populations identified plasmablasts, γδ T cells, effector memory CD4^+^ αβ T cells, NK cells, and central memory CD4^+^ αβ T cells as the most informative populations (**Fig. 5b**). Analysis of attention-weighted cell-cell communication further revealed a global reduction in intercellular signaling in RA relative to HC (see **Methods**). From these top-ranked cell types, we derived gene-attention scores and their associated signatures (**Fig. 5c**), capturing key RA-relevant mechanisms including citrullination (PADI2/4), type I interferon signaling (ISG15, IFI6), and metabolic regulation (SLC2A1, PGD). To further characterize disease-associated genes, we compared gene embeddings between conditions. Aggregating gene embeddings by condition enabled a differential analysis that identified 99 genes significantly separating RA from HC (FDR<0.05) (**Fig. 5d**). These genes formed a metabolic and structural effector module enriched for mitochondrial transcripts (MT-CO1/2/3, MT-ATP6) and matrix-remodeling components (COL6A2, PACSIN2). While this module partially overlapped with attention-derived signatures, it primarily reflected downstream effector processes rather than upstream regulatory activity. By contrast, FloREN’s attention scores highlighted regulatory hubs integrating citrullination (PADI2/4), interferon signaling (ISG15, IFI6), and complement initiation (C1QA), underscoring the complementary information captured by differential expression versus model-derived attention.

**Fig. 5.**
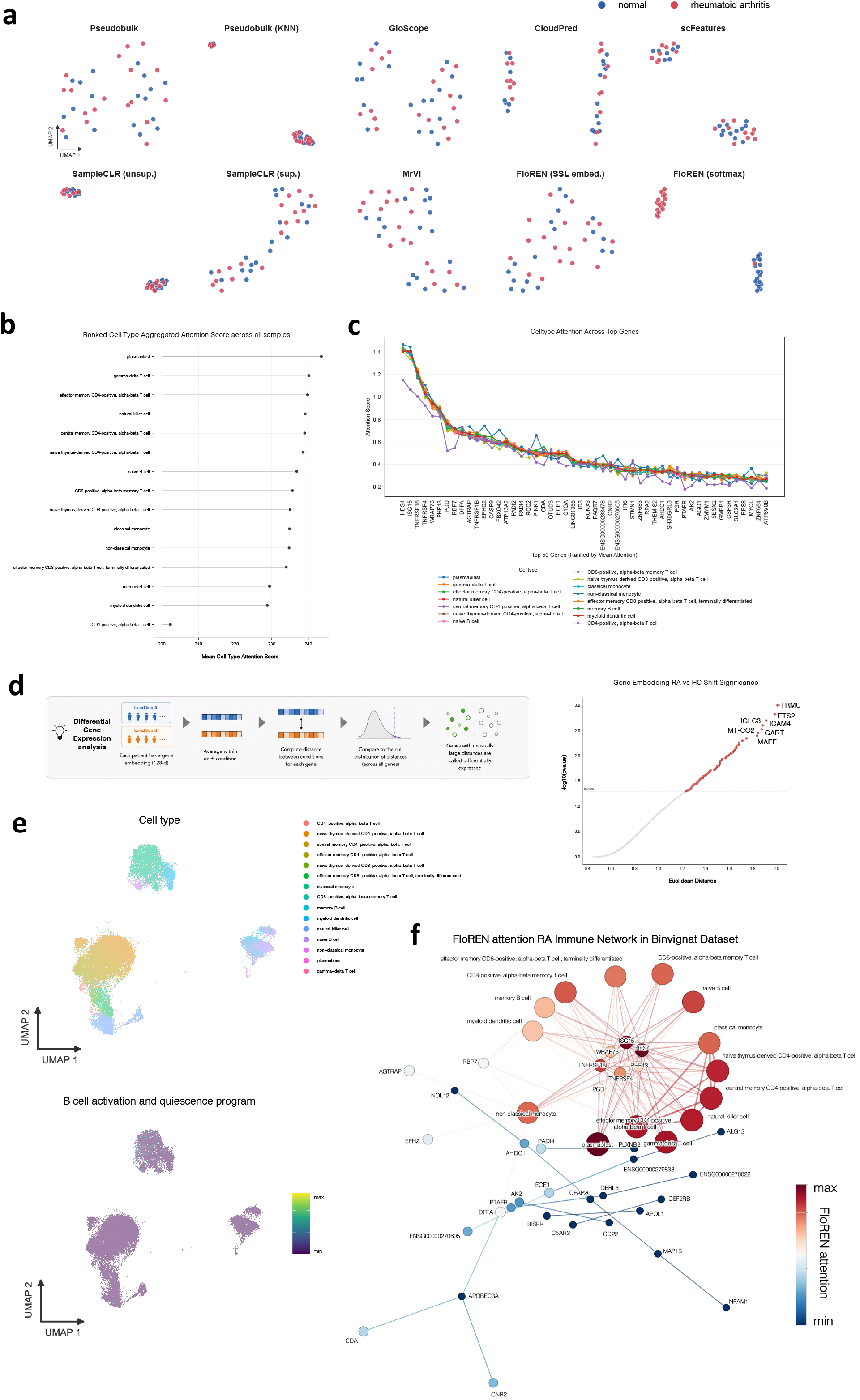
FloREN prioritizes disease-associated cell types and gene programs in rheumatoid arthritis. **a**, UMAP visualization of sample embeddings applied to the Nehar-Belaid dataset^44^ obtained from sample pseudobulk (“Pseudobulk”), kNN classifier on the pseudobulk embeddings (“Pseudobulk (KNN)”), GloScope, Cloudpred, scFeatures, unsupervised (“SampleCLR (unsup.)”) and supervised (“SampleCLR (sup.)”) SampleCLR, MrVI, FloREN’s self-supervised (“FloREN (SSL embed.)”) and final classfier (“FloREN (softmax)”). **b**, Bubble plot showing cell types ranked by their mean attention score across patients. **c**, Ranked genes according to their mean attention score across cell types. **d**, Violin plot of Euclidean distance and −log10(pvalue) showing signifcant genes on gene embeddings analysis. **e**, UMAP of cells colored by disease, cell type annotation and FloREN B cell activation and quiescence program activity. **f**, FloREN attention immune network visualization.

To place these findings in a functional context, we performed community detection on the combined set of top attention-ranked and differentially expressed genes. This identified nine gene programs, annotated via Fisher’s exact test (see **Methods**), representing coordinated immune states including interferon signaling, inflammatory monocyte priming, metabolic reprogramming, and complement activation. Notably, we observed condition-associated shifts in antigen-processing and B-cell activation programs. Increased activity of the B-cell activation program mapped specifically to memory B cells, naive B cells, and plasmablasts, refining the cell populations implicated by our earlier attention-based ranking (**Fig. 5e**). The co-regulation of FCRL1 with metabolic (GLUL) and cell-cycle (G0S2) genes further suggests a coordinated, system-wide activation state extending beyond the B-cell compartment alone.

By prioritizing key cell populations, identifying disease-relevant genes, and organizing them into coherent regulatory programs, FloREN reveals a coordinated immune network in RA (**Fig. 5f**) in which citrullination and interferon signaling act as central hubs orchestrating systemic inflammation, advancing the identification of network-level biomarkers beyond single-cell-type markers alone.

## Discussion

Single-cell RNA sequencing has transformed our understanding of cellular heterogeneity in human disease. Standard analyses typically characterize differences at the level of individual cell types between conditions, relying on expert interpretation to integrate these findings into a coherent, system-level view of the cellular environment. As single-cell atlases have grown in scale, sample representation methods have emerged as a promising strategy for summarizing this complexity into patient-level signatures. Yet even in the era of foundation models, most such methods remain either unsupervised, offering no direct link to phenotypes of interest, or supervised without interpretability, leaving the biological basis of their predictions opaque. This trade-off has confined sample representation largely to exploratory analysis, short of its potential as a tool for hypothesis-driven discovery.

FloREN (Framework for Learning Over REgulatory-Embedding Networks) addresses this gap directly. Rather than treating supervision and interpretability as competing design goals, FloREN unifies them within a single graph-based architecture: single-cell data is modeled as a heterogeneous biological network of cells and genes linked through regulatory and communicative relationships, and a combination of self-supervised and supervised objectives learns condition-relevant sample embeddings through this network via the HGTCL architecture. Because attention is computed natively over this biological graph, every prediction FloREN makes is traceable back to the specific cells, genes, and interactions that support it, turning sample representation from a black-box summarization step into a transparent, task-driven analysis in its own right.

Across five independent immunological datasets spanning RA, MS, and SLE, FloREN consistently outperformed existing sample representation approaches while preserving this interpretability at every stage of the pipeline. The framework yields three complementary outputs from a single model: supervised sample-level embeddings optimized for classification, condition-aware cell and gene embeddings that capture disease-relevant structure, and an attention network that assigns interpretable scores to nodes and interactions across the biological graph. In SLE, these sample embeddings captured not only overall disease status but also continuous disease trajectories, resolving intermediate and aggressive states consistent with independent pseudotime inference. FloREN’s cell embeddings further revealed condition-specific cellular niches, including a shift in plasmacytoid dendritic cell association consistent with their established role in type I interferon-driven autoimmunity. In MS, attention-based differential abundance analysis highlighted a distinct inflammatory and reprogramming program within monocytes, converging on regulatory hubs, including interferon-stimulated genes and complement components, that mirrored mechanisms independently identified in SLE and RA. In RA, gene embeddings enabled differential expression analysis alongside attention-based prioritization of a B-cell activation program centered on plasmablasts, which emerged as a core hub within a broader systemic inflammation network. Notably, the recurrence of type I interferon and complement-related signatures across all three, biologically and etiologically distinct diseases suggests that FloREN is not merely fitting dataset-specific noise but recovering a shared regulatory architecture underlying immune-mediated inflammation, precisely the kind of network-level convergence that motivated our use of IMIDs as a stringent evaluation setting.

These results illustrate the central contribution of this work: FloREN demonstrates that supervision and interpretability need not be in tension in sample representation learning. Where prior unsupervised methods offer interpretability without a clear predictive objective, and prior supervised methods offer strong predictive performance with limited mechanistic insight, FloREN’s graph-native attention mechanism ties both together. Every embedding dimension and every attention score is grounded in an explicit biological entity or interaction, rather than an abstract latent coordinate. In this sense, FloREN aims to mark a step forward for sample representation methods: moving the field toward approaches that combine a clearly defined, task-relevant learning objective with biologically interpretable network modeling, rather than requiring a trade-off between the two.

Despite its strengths, several limitations should be considered. First, FloREN’s computational complexity scales with both the number of samples and the density of graph edges, making training on large-scale datasets resource intensive. Future work could mitigate this through graph sparsification, subgraph sampling, or more efficient mini-batch training strategies for heterogeneous graphs. Incorporating pretrained or foundation models to initialize cell and gene embeddings represents a promising direction to further improve performance and scalability. Second, the framework might rely on curated prior knowledge (e.g., gene-gene interactions and cell-cell communication databases) and preprocessing choices such as gene selection and cell type annotation. Variability or bias in these inputs may influence downstream representations and interpretability. Applications outside the immune system will require new prior knowledge generation if it wants to be used. Third, while attention scores highlight statistical importance within the model, they do not by themselves establish causal or mechanistic roles; complementary perturbation-based validation will be necessary to confirm the functional relevance of prioritized genes and populations, such as CHI3L2 in MS or the FCRL1-centered program in RA. Finally, our evaluation focused on IMIDs, chosen for their network-level pathology; extending FloREN to other disease contexts, including solid tumors and neurodegenerative disease, will be important to establish the generality of this approach.

Looking forward, FloREN’s modular graph construction makes it well suited to increasingly complex experimental designs. Extensions to longitudinal cohorts could enable direct modeling of temporal disease dynamics rather than the cross-sectional trajectory inference used here, while integration with spatial transcriptomics or additional omics layers may yield richer, tissue-resolved representations of cellular organization and regulation.

In summary, FloREN establishes a supervised, interpretable framework for sample representation in single-cell analysis. By uniting biologically informed graph modeling with attention-based interpretability, FloREN reframes sample representation as a mechanistically grounded discovery task, enabling scalable identification of the cellular programs and regulatory networks that underlie complex human diseases.

## Data and code availability

This paper analyzes existing, publicly available data, accessible at GEO and CELLxGENE through the accession numbers in Methods.

All code, installation instructions, and usage tutorials for FloREN are publicly available at https://github.com/iclemente99/FloREN.

## Acknowledgements

This work was supported by the European Union’s Horizon Europe research and innovation programme. We thank Dr. Yvan Saeys (VIB-UGent Center for Inflammation Research) and Dr. Mikel Hernaez (CIMA, University of Navarra) for their valuable input on framework construction and validation. We also acknowledge the 3TR (grant agreement No 831434) and PRECISESADS (grant agreement No 115565) consortium, which guaranteed the data used in the prior knowledge gene associations. We also thank Dr. Alexandra-Chloé Villani and Dr. Gary Reynolds and all the Villani-Reynolds Lab (Massachusetts General Hospital, Broad Institute Harvard-MIT and Harvard Medical School), especially Dr. Sergio Aguilar-Fernandez, for their continues feedback and suggestions for improvements. We finally want to also thank Dr. Francesco Craighero at Holger Heyn’s lab in Centro Nacional de Análisis Genómico for the machine learning feedback recieved. Some figures were created with BioRender.com.

## Author contributions

ICL, CJ and NF designed the study. ICL developed FloREN, performed the analyses and drafted the manuscript. ICL, CJ and NF participated in the writing of the article, SH and DC revised it intellectually critically, and all authors approved the submitted version.

## Declaration of generative AI and AI-assisted technologies

During the preparation of this work, the author(s) used ChatGPT and Gemini in order to improve English writing and clarify figures. After using these tools, the author(s) reviewed and edited the content as needed and take(s) full responsibility for the content of the publication.

## Methods

### Data collection

Data used was collected from public databases: GSE135779, GSE138266, GSE133028, GSE163005, GSE174188 and https://cellxgene.cziscience.com/collections/e1a9ca56-f2ee-435d-980a-4f49ab7a952b.

### Data preprocessing and normalization

Datasets GSE174188 and https://cellxgene.cziscience.com/collections/e1a9ca56-f2ee-435d-980a-4f49ab7a952b were already preprocessed. Databases not preprocessed were loaded into a *Seurat* R package^60^ object using ensemble id gene names. All of them were filtered under the same conditions: Number of unique genes per cell between 200 and 2500, and mitochondrial RNA below 10 percent. Percentage of mitochondrial RNA filtering ensures that cells sequenced are living healthy cells and was calculated with the collection of human mitochondrial RNA from *scCustomize* package^61^. Then, datasets were processed independently following the standard Seurat protocol^62^ with log-normalization and scale it to allow further analysis on them. Transformation of genes to TFs was done with *DecoupleR* R package^63^ trough Area Under the recovery curve for each Cell (AUCell) score.

The Binvignat dataset^48^ includes peripheral blood mononuclear cell (PBMC) samples from 18 rheumatoid arthritis (RA) patients and 18 matched healthy controls. For each sample, we built a heterogeneous graph using the 2000 most variable genes. Cell-gene edges were inferred from expression counts, gene-gene edges from known interactions, and cell-cell connections from communication scores. The Straeten dataset^49^ contains cerebrospinal fluid (CSF) samples from 17 multiple sclerosis (MS) patients and 11 controls. Graphs were constructed using variable genes and included cell-gene and gene-gene interactions. The Schafflick dataset^50^ extends this design by incorporating both PBMC and CSF samples (6 MS and 5 controls per compartment), and we used all preprocessed genes to assess FloREN’s capacity to handle broader expression variability. The Nehar-Belaid dataset^51^ includes 40 SLE patients (7 adults and 33 children) and 16 healthy controls. To emphasize regulatory context, we converted the gene matrix to TF activity profiles and constructed graphs with TF-cell and TF-TF edges. The Perez dataset^52^ consists of 162 SLE and 99 control PBMC samples. Due to its larger scale, we optimized computational efficiency by using only TFs derived from the 2000 most variable genes, constructing TF-cell and TF-TF graphs per patient.

### Autoencoder (AE)

The Autoencoder (AE) layer serves to unify representations of cells and genes, producing 256-dimensional embeddings for both entities. These embeddings provide a well-defined node representation when constructing graphs.

The input of the Autoencoder is the matrix *X*^*G*^ = {*i* = 1,2, … *I*; *j* = 1,2, … *J*} where I denote the number of genes and J the number of cells. Each element *x*_*ij*_ represents the normalized expression level of gene *i* and cell *j* from single-cell RNA sequencing (scRNA-seq) data.

We implement two separate autoencoders to generate initial embeddings:

- **Cell Autoencoder:** Reduces the gene-dimensional representation of each cell from *I* to 512 and then to 256 dimensions.
- **Gene Autoencoder:** Reduces the cell-dimensional representation of each gene from *J* to 512 and then to 256 dimensions.

Thus, every cell and gene are assigned an initial 256-dimensional representation. The number of reduced dimensions is optimized as a hyperparameter, varying based on dataset characteristics. The final output is a reconstructed matrix 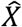 that retains the same dimensions as the original *X*.

The loss function for the cell autoencoder is defined as the Mean Squared Error (MSE) between *X* and its reconstruction 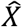:

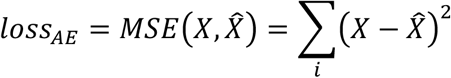

Similarly, the gene autoencoder extracts low-dimensional gene features across all cells. It follows the same structure as the cell autoencoder but operates on the transposed input matrix *X*^*T*^.The corresponding loss function is:

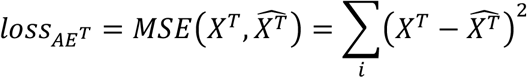

By constructing cell and gene embeddings independently, this autoencoder-based representation enhances graph structure robustness when integrated into downstream tasks.

### Cell-feature graph construction

A graph is a mathematical structure represented as *G* = (*V, E*), where *V* is the set of nodes (the elements that form the graph) and *E* the set of edges connecting them. In this work, we construct a heterogeneous graph, incorporating various node and edge types to capture biological interactions comprehensively.

- **Heterogeneous graph**: A graph containing multiple types of nodes and/or edges. We define it as *G* = (*V, E, A, R*), where we have the *V* represents nodes, *E* represents edges, *A* represents the node type union, and *R* represents the edge type union.
- **Node and edge type mapping functions**: We define *τ*(*v*): *V* → *A* and *ϕ*(*e*): *E* → *R* as the mapping function for node types and edge types, respectively.
- **Node meta relation:** For a node pair of *v*_1_ and *v*_2_ linked by an edge *e*_1,2_, the meta relation between *v*_*i*_ and *v*_*j*_ is denoted as ⟨*τ*(*v*_*i*_), *ϕ*6*e*_*ij*_7, *τ*6*v*_*j*_7⟩.

Our approach constructs an Undirected Binary Heterogeneous Graph with at least two hierarchical levels: cells and features (can be genes or TFs). The methodology is extendable to more complex structures, such as cells-TFs-genes interactions.

#### Gene-cell connections

We define a bipartite gene-cell heterogeneous graph based on the input expression matrix *X*^*G*^ = {*i* = 1,2, … *I*; *j* = 1,2, … *J*}, where *I* is the number of genes and *J* is the number of cells and each *x*_*ij*_ represents the normalized expression of gene *i* with cell *j*. The graph consists of two node types (genes and cells) and one edge type (gene-cell). Nodes are defined as: *V* = *V*^*C*^ ∪ *V*^*G*^ where *V*^*G*^ = {*i* = 1,2, … *I*} denotes all genes, and *V*^*C*^ = {*j* = 1,2, … *J*} denotes all cells. Edges *E* = {*e*_*ij*_} exist if *x*_*ij*_ > 0, assigned weight *ω*(*e*_*ij*_) = 1, otherwise, *ω*(*e*_*ij*_)= 0.

#### Tf-cell connections

Using the TF activity matrix *X*^*tf*^, derived from *DecoupleR* R package^63, we construct a TF-cell bipartite graph with TFs and cells as node types and TF-cell as edges. The^graph is defined as: *V* = *V*^*C*^ ∪ *V*^*tf*^, where *V*^*tf*^ = {*i* = 1,2, … *I*} denotes all TFs, and *V*^*C*^ = {*j* = 1,2, … *J*} denotes all cells. Edges *E* = {*e*_*ij*_} exist if *x*_*ij*_ > 0, assigned weight *ω*(*e*_*ij*_) = 1, otherwise, *ω*(*e*_*ij*_) = 0.

#### Gene Regulatory Network (GRN) inference

For gene-gene relationships, we construct a homogeneous gene-gene graph based on the input matrix *X*^*G*^ = {*i* = 1,2, … *I*; *j* = 1,2, … *J*} with I genes and J cells. The gene-gene graph is defined as *V* = *V*^*F*^, where *V*^*F*^ = {*i* = 1,2, … *I*}, and *E* = {*e*_*ij*_} represents gene-gene associations. Edges are assigned using a custom-designed score:

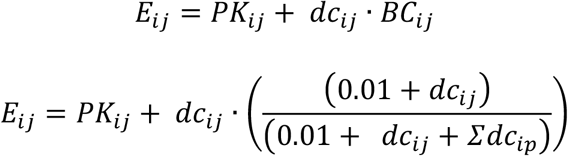

Where *PK*_*ij*_ represents prior biological knowledge, obtained via STRING (*stringdb* Python module^64^) and INNATEDB (*PSIQUIC* REST API), *dc*_*ij*_ can be classical linear Pearson correlation or the distance correlation capturing both linear and nonlinear associations with *dcor* R package^65^and the final term is a Bayesian probability, estimating the likelihood that an observed data-driven association is biologically causal. The constant 0.01 is introduced for numerical stability (More details in another section).

An explicit threshold was applied to filter gene-gene associations. Specifically, edges were retained only when the score satisfied *E*_*ij*_ ≤ 0.7; in such cases, the weight of the corresponding edge was set to *ω*(*e*_*ij*_) = 1, otherwise, *ω*(*e*_*ij*_) = 0. For TF-TF graphs, edges were computed analogously, by aggregating gene-level connections for the transcription factors they regulate.

#### Cell-cell connections

To capture cellular communication, we construct a cell-cell homogeneous graph from the matrix *X*^*C*^ = {*i* = 1,2, … *J*; *j* = 1,2, … *J*} with J cells derived using the *CellChat* R package (version 1.6.0)^66^. The cell-cell graph is defined as *V* = *V*^*C*^, where *V*^*C*^ = {*i* = 1,2, … *J*} denotes all cells, and *E* = {*e*_*ij*_} represents cell-cell interactions.

*CellChat* infers communication probabilities based on: (1) Ligand-receptor interactions from a curated database, (2) average ligand expression in one cell type and receptor expression in another and (3) heteromeric complexes and cell-type proportions.

Again, arbitrary threshold can be used to filter cell-cell associations. For example, if *x*_*ij*_ ∈ *X*^*C*^ > 0.5, the weight of the corresponding edge *ω*(*e*_*ij*_) = 1, otherwise, *ω*(*e*_*ij*_) = 0.

### Single Cell Graph Contrastive Learning

We introduce a self-supervised architecture designed to discern specific patterns within the informative Heterogenous Graphs applied to single-cell networks. Building upon the Graph Contrastive Learning (*GraphCL*) framework^67^, we modified the GNN-based encoder with an HGT-based encoder and incorporate a classification loss function to enhance performance. Recognizing the limitations observed when employing GSSL solely for pretraining followed by secondary multi-task learning finetuning, we have restructured the standard architecture to address these challenges (More details in another section).

In this modified approach, pretraining is conducted by maximizing the agreement between two augmented views of the same graph using a contrastive loss in the latent space. Here, an augmented view refers to a transformed version of the input graph, generated through operations such as randomly dropping nodes, masking node features or dropping edges, while preserving the overall biological semantics. Additionally, a Multi-Layer Perceptron (MLP) is employed to classify the latent representations via a classification loss. The framework comprises six key components:

#### Graph data augmentation

This process generates novel and semantically consistent data by applying specific transformations that preserve the original semantic labels. Each graph *G* undergoes graph data augmentations to obtain two correlated views *Ĝ*_*i*_, *Ĝ*_*j*_. In the results section we only used node dropping augmentations.

#### HGT-based encoder

A HGT encoder, referred to as *HGT*_*loyer*_ extracts graph-level representation vectors *h*_*i*_, *h*_*j*_ from the augmented graphs *Ĝ*_*i*_, *Ĝ*_*j*_. In our implementation, this encoder is a one-layer HGT with no activation function. The representation *h*_*i*_ is computed as the mean of the output nodes from *HGT*_*i*_ (*Ĝ*_*i*_). The dimensionality of *h*_*i*_, *h*_*j*_ is treated as a hyperparameter, optimized based on the dataset and training objectives; in our standard approach, we set this dimension to 128.

#### MLP classifier

A four-layer MLP with GeLU activation functions is applied to the mean of *h*_*i*_ and *h*_*j*_, producing an output vector of size *n*, where *n* corresponds to the number of classification labels. The architecture of this MLP can be adapted to suit specific goals and datasets.

#### Contrastive loss function

The contrastive loss function *L*_*con*_ aims to maximize the consistency between positive pairs *h*_*i*_, *h*_*j*_ while distinguishing them from negative pairs. We employ the normalized temperature-scaled cross-entropy loss (NTXent) for this purpose^68–69^.

#### Classification loss function

The classification loss function *L*_*class*_ to ensure the learning of condition-based patterns from the mean of the graph-level representation vectors *h*_*i*_, *h*_*j*_, we utilize the cross-entropy loss (CE).

#### Dynamic Multitask Learning

We implement a dynamic multitask learning approach by adjusting the weighted loss, defined as:

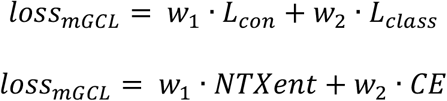

where *w*_1_ and *w*_2_ are weights that balance the contributions of the contrastive and classification loss components, respectively.

Thus, the *loss*_*GraphCL*_ *n*th graph is defined as:

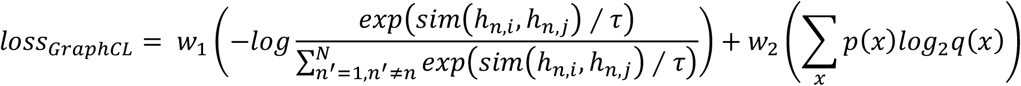

Where, *sim* (*h*_*n,i*_, *h*_*n,j*_) = *h*_*n,i*_^*T*^*h*_*n,j*_ / ‖*h*_*n,i*_‖‖*h*_*n,j*_‖, denotes the cosine similarity function, calculated as:

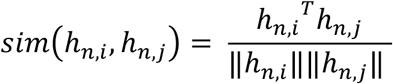

The parameter τ represents the temperature parameter, and *x* defines the different label outcomes for the patients.

For each patient graph, we generate two augmented versions and apply the HGT-based encoder to both, extracting representation vectors. The contrastive loss is applied to these vectors, and their mean is fed into the MLP classifier to compute the classification loss.

### Heterogeneous Graph Transformer

We propose the HGT model^71^ for the GNN-encoder head. The HGT is employed to learn expressive embeddings for both genes and cells, and to estimate their mutual influence via attention mechanisms. Its input is the augmented heterogenous graphs. Its output is refined embeddings and attention scores that reflect biological relevance.

- **Target node and source node:** In the context of HGT, a target node *v*_*t*_ is the node currently being updated, while a source node *v*_*s*_ is any neighbor connected to it via an edge *e*_*s,t*_. These roles are crucial for directed attention and message passing.
- **Neighborhood graph of target node**: For each *v*_*t*_, a local subgraph *G*′ = (*V*′, *E*′, *A*′, *R*′) is created, which includes the target, its neighbors *N*(*v*_*t*_), and associated node types *A*′ and edge types *R*′. This local structure allows the transformer to focus on relevant context.

#### Multi-head attention mechanism and linear mapping of vectors

Each HGT *l*^*th*^ layer uses multi-head attention to learn relationships across different embedding subspaces. The embedding of *v*_*t*_ and *v*_*s*_ on the *l*^*th*^ layer are denoted as *H*^*l*^ [*v*_*t*_] and *H*^*l*^ [*v*_*s*_]. Each head *h* learns:

- A query vector: 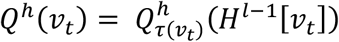
- A key vector: 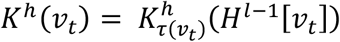
- A value vector: 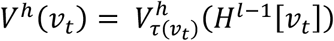

Each node type has its own linear projection layers to adapt to structural heterogeneity.

#### Heterogeneous mutual attention

The mutual attention between *v*_*t*_ and *v*_*s*_ is calculated with a specialized attention operator:

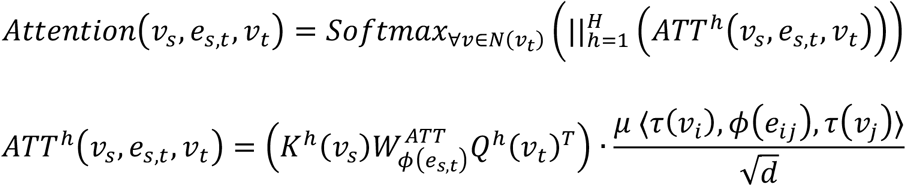

Here, *μ* is a learnable scalar denoting the importance of the meta-relation (node-type, edge-type, node-type).

#### Heterogenous message passing

Each source node passes a message to the target node:

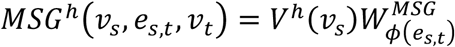

These are concatenated across all heads:

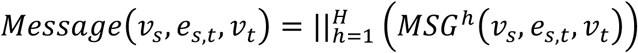

#### Target specific aggregation

The new embedding of *v*_*t*_ at *l*^*th*^ layer is obtained by aggregating all messages weighted by attention:

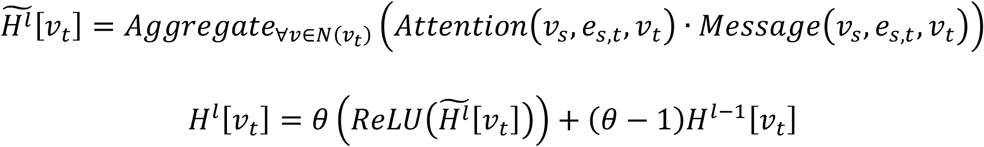

θ is a trainable parameter controlling residual blending.

#### Determination of edges attention

After the final HGT layer, we obtain edges attention scores:

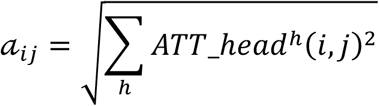

These values reflect how important a node *i* is to the identity of node *j*, which can later inform downstream biological interpretation.

#### HGT training on subgraph

As HGT trains on how to weight the different types of edges. HGT can train on the augmented subgraphs and apply the learned weights to the full graph.

### Sample Representations Evaluation Settings

The SPARE metrics (Sample Preservation and Representation Evaluation)^53^ are a set of three complementary scores designed to quantitatively assess how well a low-dimensional sample embedding preserves biologically meaningful structure while removing unwanted technical variation.

**Information retention & batch effect removal** measures how strongly relevant biological/clinical covariates (e.g., disease status) are preserved in the embedding and how completely technical covariates (e.g., batch) are removed. It uses k=3 nearest-neighbor classification/regression to predict each sample’s covariate value from its neighbors in embedding space, then computes a corrected macro-F1 score (clipped at 0) for categorical variables or absolute Spearman correlation for continuous/ordinal ones; relevant scores are averaged directly, while technical (batch) scores are inverted (1 – retention) so higher values mean better removal.

**Biological trajectory preservation** evaluates whether continuous processes (e.g., age) are correctly ordered in the embedding. It computes diffusion pseudotime starting from the sample with the minimal value of the trajectory covariate (assumed earliest point), then reports the absolute Spearman correlation between this pseudotime and the true covariate value.

**Replicate robustness** checks whether biologically replicate samples (e.g., multiple samples from the same patient/individual) are placed closer to each other in the embedding than to non-replicates. For each replicate pair, it calculates the fraction of all other samples that are farther from one replicate than its pair partner is, averages over both directions of the pair, and finally averages across all such pairs.

These three components are combined into a single aggregated SPARE score using a weighted average (information retention and trajectory preservation at weight 1, replicate robustness at weight 1, batch removal at weight 0.5 to avoid biasing toward random-like embeddings), normalized by the total weight (3.5) to produce a final value in [0,1], where higher is better.

For the results, we used Disease as relevant information retention, we used Batch for irrelevant information removal, we used Age for trajectory conservation, and we used Sex replicate robustness.

### Predictive Evaluation Settings

To assess the predictive performance of sample representation methods, we benchmarked based on 5-fold cross-validation kNN classifier performed on the output sample representation embeddings of each method. We calculated the F1 score, Area Under the Receiver Operating Characteristic Curve (AUC), and Balanced Accuracy (BalAcc). All metrics were calculated using *sklearn* Python module^72^.

### Clustering Statistics on Sample Embeddings

Patients level spaces were also evaluated with both unsupervised and supervised clustering scores. We used the following unsupervised clustering score: Calinski-Harabasz Index (CH), Average Silhouette Score (AWS) and Davies-Bouldin Index (DB). On the other hand, we used also three other supervised clustering score: Adjusted Rank Index (ARI) and Adjusted Mutual Information (AMI). All metrics were calculated using *sklearn* Python module^72^.

Clustering for the supervised metrics was performed using the *KMeans* function from the *sklearn* module in Python^72^. To infer trajectories within the learned patient spaces, we used the *slingshot* package from R^55^.

### Cell niche analysis

FloREN generates a 128-dimensional embedding vector for each cell, representing its learned biological state within the heterogeneous graph. To obtain condition-specific representations, cell embeddings were first grouped by annotated cell type and averaged across all cells belonging to that cell type within each patient. Subsequently, patient-level cell type embeddings were averaged across all patients from the same condition, yielding a single representative embedding for each cell type in each condition.

To quantify relationships between cell types, we computed the Euclidean distance between all pairs of condition-specific cell type embeddings,

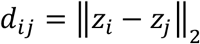

Where *z*_*i*_ and *z*_*j*_ denote the mean embedding vectors of cell types i and j, respectively. The resulting pairwise distance matrix was visualized by hierarchical clustering using average linkage.

The underlying assumption is that FloREN embeds cell types participating in coordinated biological programs into nearby regions of the latent space. Consequently, cell types with smaller Euclidean distances are interpreted as belonging to the same functional cellular niche, whereas increased distances indicate distinct or condition-specific cellular states. Comparing the distance matrices between conditions therefore enables the identification of niche reorganization associated with disease.

### Differential Abundance Analysis

The FloREN attention network assigns an attention score to each edge in the heterogeneous graph constructed for every patient. For each node (cell) in the graph, a node-level attention score was obtained by aggregating the attention scores of all edges connected to that node. Formally, if *e*_*ij*_ denotes the edge between nodes *i* and *j* with attention weight *a*_*ij*_, the attention score for node *i* is computed as:

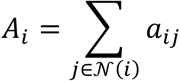

where *N*(*i*) is the set of neighboring nodes connected to node *i*.

Once a cell-level attention score 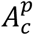 was obtained for every cell *c* in patient *p*, the total attention score for each cell type *t* in patient *p* was calculated by summing the attention scores of all cells belonging to that cell type:

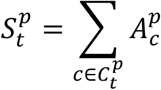

where 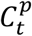 represents the set of all cells of type *t* in patient *p*.

Under the null hypothesis that a given cell type *t* has the same biological importance (as captured by the attention mechanism) across two conditions (e.g., disease vs. control), the total attention sums 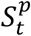 are expected to follow the same distribution in both groups. A significant difference in the distribution of 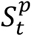 between the two conditions indicates differential abundance or differential importance of that cell type.

To test this hypothesis, a non-parametric two-sided Mann-Whitney U test was performed to compare the total FloREN attention sums 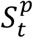 for each cell type *t* between the two patient groups. The resulting p-values were adjusted for multiple testing using the Benjamini-Hochberg false discovery rate (FDR) correction. Cell types with an FDR-adjusted p-value below a predefined significance threshold (typically 0.05) were considered differentially abundant between the conditions.

### Differential Gene Expression Analysis

The FloREN model produces a 128-dimensional gene embedding vector for each gene in every patient. To obtain a condition-specific representation, the gene embeddings were first averaged across all patients within each condition. Formally, for a given gene *t*, the condition-specific embedding for condition *c* is defined as:

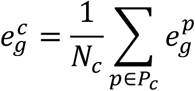

where 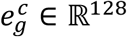 is the embedding vector of gene *g* in patient *p, P* is the set of all patients belonging to condition *c*, and *N*_*c*_ = |*P*_*c*_| is the number of patients in that condition.

Under the assumption that these embeddings capture condition-aware biological information, the difference between the embeddings of the same gene across the two conditions reflects differential gene relevance or activity. For each gene g, the distance between its two condition-specific embeddings was computed as:

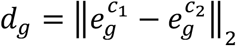

where ‖·‖_2_ denotes the Euclidean (L2) norm.

To assess statistical significance, an empirical p-value was calculated for each gene based on the null distribution of distances across all genes. Specifically, the p-value represents the probability of observing a distance at least as large as *d*_*K*_ under the null hypothesis that the observed distance is due to random variation. The empirical p-value is given by:

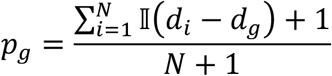

where *d*_*i*_ is the distance for the *i*-th gene, *N* is the total number of genes, and I(·) is the indicator function. This formulation corresponds to the standard rank-based empirical p-value with continuity correction (adding 1 to both numerator and denominator to avoid p = 0).

Genes with a small empirical p-value indicate a significantly larger embedding distance between conditions compared to the background distribution of all genes, and were therefore considered differentially expressed. This approach provides a differential gene expression analysis that is robust to technical noise such as batch effects, as the condition-aware embeddings are learned directly from the graph attention mechanism.

### Cell-cell communication profiling

FloREN learns an attention score for every edge in the heterogeneous graph of each patient, including cell-cell, gene-cell, and gene-gene interactions. To characterize cell-cell communication, we isolated the attention scores corresponding to cell-cell edges. Using the cell type annotations, we then aggregated the attention scores of all interactions between each pair of annotated cell types, yielding a cell type-cell type interaction matrix for every patient.

The resulting matrix represents the relative importance assigned by FloREN to communication between cell populations, where higher attention scores indicate interactions that contribute more strongly to the learned disease representation. Finally, interaction matrices were averaged across patients within each condition to generate condition-specific cell-cell communication profiles, enabling qualitative comparison of communication patterns between disease states.

### Unique Cell Type Gene Signature Decomposition

To derive cell type-specific gene signatures, we adapted the gene program scoring strategy described by Hachonen et al.^73^. Starting from FloREN’s attention-weighted edge matrix (edges defined as ⟨source, target, attention⟩), we first isolated edges involving cell type-specific nodes. Gene-level attention scores were then aggregated per cell type, resulting in a matrix of dimensions (cell types × genes), where each entry represents the aggregated attention weight of a gene within a given cell type. This matrix is conceptually analogous to the (gene programs × genes) matrix obtained from non-negative matrix factorization (NMF) in the original framework.

To characterize cell type-specific gene signatures, we ranked genes within each cell type using a scaled weighting scheme designed to prioritize both high relevance and uniqueness. For gene *i* and cell type *j*, the scaled weight *WS*_*ij*_ was defined as:

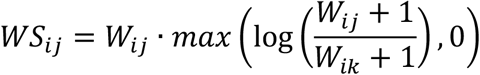

Where *W*_*ij*_ denotes the aggregated attention weight of gene *i* in cell type *j*, and *W*_*ik*_ is the maximum aggregated weight of gene *i* across all other cell types *k* ≠ *j*. This formulation emphasizes genes with high attention within a given cell type (first term) while penalizing genes that are broadly shared across cell types (second term).

We calculated the cumulative sum of *WS*_*ij*_ and select the genes that explain 99% of each Gene Program. This way we obtained of around 250 genes that are unique for each Gene Program.

### Gene program calculation

To identify disease-associated gene programs, we leveraged FloREN’s attention scores to construct condition-specific gene regulatory networks. For each dataset, we first selected the top five cell types based on their aggregated cell-level attention scores. Within these cell types, we further selected the top 250 genes ranked by gene-level attention.

Using the attention-weighted gene-gene edges, we extracted gene regulatory networks (GRNs) for each patient, restricted to the selected genes. These patient-level GRNs were then aggregated by disease condition, resulting in one condition-specific GRN per group. In these networks, both nodes and edges were weighted according to their corresponding FloREN attention scores, thereby preserving the relative importance learned by the model.

To identify coherent gene programs, we applied the Leiden community detection algorithm^74^ to each condition specific GRN. Gene programs were defined as communities containing at least four genes. Using this approach, we identified six rheumatoid arthritis (RA) gene programs and seven healthy donor (HD) gene programs.

### Annotation target enrichment in gene expression programs

Transcription factor targets, immune hallmark gene sets, and pathway annotations were obtained from the Molecular Signatures Database (MSigDB; http://www.gsea-msigdb.org/gsea/msigdb/, accessed February 2026)^77^. For each gene program, enrichment of these target sets was assessed using a Fisher’s exact test applied to the overlap between the program’s top-ranked genes and each reference gene set.

A target set was considered significantly associated with a given gene program if it satisfied both criteria: (i) a false discovery rate (FDR) below 0.1 after multiple testing correction, and (ii) a minimum overlap of three genes between the gene program and the target set.

